# *Wolbachia* endosymbionts subvert the endoplasmic reticulum to acquire host membranes without triggering ER stress

**DOI:** 10.1101/448365

**Authors:** Nour Fattouh, Chantal Cazevieille, Frédéric Landmann

## Abstract

The reproductive parasite *Wolbachia* are the most common endosymbionts on earth, present in a plethora of arthropod species. They have been introduced into mosquitos to successfully prevent the spread of vector-borne diseases, yet the strategies of host cell subversion underlying their obligate intracellular lifestyle remain to be explored in depth in order to gain insights into the mechanisms of pathogen-blocking. Like some other intracellular bacteria, *Wolbachia* reside in a host-derived vacuole in order to replicate and escape the immune surveillance. Using here the pathogen-blocking *Wolbachia* strain from *Drosophila melanogaster,* introduced into two different *Drosophila* cell lines, we show that *Wolbachia* subvert the endoplasmic reticulum to acquire their vacuolar membrane and colonize the host cell at high density. *Wolbachia* redistribute the endoplasmic reticulum to increase contact sites, and time lapse experiments reveal tight coupled dynamics suggesting important signalling events or nutrient uptake. They however do not affect the tubular or cisternal morphologies. A fraction of endoplasmic reticulum becomes clustered, allowing the endosymbionts to reside in between the endoplasmic reticulum and the Golgi apparatus, possibly modulating the traffic between these two organelles. Gene expression analyses and immunostaining studies suggest that *Wolbachia* achieve persistent infections at very high titers without triggering endoplasmic reticulum stress or enhanced ERAD-driven proteolysis, suggesting that amino acid salvage is achieved through modulation of other signalling pathways.

**Author summary:** *Wolbachia* are a genus of intracellular bacteria living in symbiosis with millions of arthropod species. They have the ability to block the transmission of arboviruses when introduced into mosquito vectors, by interfering with the cellular resources exploited by these viruses. Despite the biomedical interest of this symbiosis, little is known about the mechanisms by which *Wolbachia* survive and replicate in the host cell. We show here that the membrane composing the *Wolbachia* vacuole is acquired from the endoplasmic reticulum, a central organelle required for protein and lipid synthesis, and from which originates a vesicular trafficking toward the Golgi apparatus and the secretory pathway. *Wolbachia* modify the distribution of this organelle to increase their interactions with this source of membrane and likely of nutrients as well. In contrast to some intracellular pathogenic bacteria, the effect of *Wolbachia* on the cell homeostasis does not induce a stress on the endoplasmic reticulum. One of the consequences of such a stress would be an increased proteolysis used to relieve the cell from an excess of misfolded proteins. Incidentally, this shows that *Wolbachia* do not acquire amino acids from the host cell through this strategy.

## Introduction

The alpha-proteobacteria *Wolbachia-Wb-* are the most common endosymbionts encountered in nature, present in a plethora of terrestrial arthropod hosts, and in filarial nematode species. These reproductive parasites have developed a wide range of symbiotic interactions, from facultative to mutualistic [1]. In all instances, they are vertically transmitted through the female germline but also colonize the soma [2]. The tissues that are infected can differ from one host species to another, as well as the *Wolbachia* intracellular titer. Although the highest titers are often observed in the germline, they vary considerably among wild isolates of specimens within a single species [3]. While *Wolbachia* intrinsic factors can be responsible for targeting specific cell types acting as reservoirs, i.e. the somatic stem cell niche in the *Drosophila* ovary [4], they can also influence the degree of intracellular replication. Such is the case for the pathogenic *Wolbachia* strain wMelpop, that possesses a region of eight genes called octomom, whose degree of amplification dictates the bacterial titer and the virulence [5]. Conversely, the host genetic background also exerts a profound influence on the bacterial ability to replicate. When the wMel strain naturally hosted in the fruit fly *Drosophila melanogaster* is transferred into the closely related *Drosophila simulans* species, mature oocytes appear dramatically more infected [6]. Therefore, depending on the permissivity of the genetic background, different cell types can harbor a wide range of endosymbiontic titers. As a consequence, the impact of a given *Wolbachia* strain on the cellular homeostasis, and the degree of subversion exerted on organelles to satisfy their obligate intracellular lifestyle can potentially induce variable phenotypes, i.e. in terms of nutrient demand, stress or cell innate immune responses.

These past years have seen a resurgence of interests in *Wolbachia* because they can be a drug target to fight parasitic filarial diseases [7], and because of their ability to compromise transmission of vector-borne arboviruses [8]. In the latter case, the wMel strain has been favored and introduced into mosquito vectors because it does not induce a fitness cost [9,10], allowing a spread through wild populations of mosquitos. Although the mechanisms by which *Wolbachia* block the pathogen transmission are not fully understood, a clearer picture starts to emerge. However among recent studies, somewhat contradictory results have been reported, reflecting a variety of phenotypes under environmental influence (for a review see [11]). Typically, the role of *Wolbachia-induced* innate immunity priming in pathogen interference is still an object of debate, although viral replication inhibition can be achieved by wMel without inducing an upregulated expression of anti-microbial peptide genes [12,13]. *Wolbachia* depend on host nutrients such as amino acids and lipids [14,15], but they potentially provision their hosts to act in some instances as nutritional symbionts. Hence, the cost and benefit associated with a *Wolbachia* infection are certainly variable. Nonetheless their intracellular lifestyle involves a competition with viruses for subverting the same limited resources. Cholesterol and lipid homeostasis are modulated in the presence of *Wolbachia* [16] and account for their pathogen-blocking effect, limiting the viral access to these metabolites essential to their replication [17,18]. If a persistent infection with *Wolbachia* endosymbionts exerts a cellular stress, it should not affect the host viability. An Endoplasmic Reticulum-ER-stress response has been described to be associated with *Wolbachia* [18,19]. The ER is involved in lipid metabolism, protein synthesis and their proper folding as well as post-translational modifications, and is the source of vesicular trafficking with the Golgi apparatus [20]. Because of its central role in the host cell metabolism, the ER is often subverted by viruses and intracellular bacteria [21,22]. When the cell homeostasis is perturbed to the point that misfolded proteins accumulate in the ER, an Unfolded Protein Response-UPR-is triggered. In order to restore homeostasis, the ER protein folding capacity is increased through chaperone release in the ER lumen and upregulation of chaperone and UPR sensor genes; translation is reduced; and an ER-associated degradation ERAD-pathway is upregulated. If the stress is prolonged, cell dysfunctions occur and cell death is eventually induced [23]. Accordingly, some intracellular bacteria have learned to subvert and control the UPR to avoid such fate [22]. It is therefore intriguing that an ER stress has been reported or suggested by some studies and up to date invoked as a consequence of a *Wolbachia* infection. More specifically, proteomic studies suggest a mild upregulation of some UPR related genes, although it should be noted that they were carried out with the life-shortening pathogenic strain wMelpop [18]. A recent study using RNAi screening in *Drosophila* cells coupled to electron microscopy observations, highlights the requirement of an ERAD ubiquitin ligase to maintain a normal *Wolbachia* titer, and reports a close subcellular vicinity between *Wolbachia* and a morphologically aberrant ER [19]. This study suggests that an ERAD-derived proteolysis is induced by *Wolbachia* to salvage amino acids. In the present study, we seek to clarify the link between *Wolbachia* and the ER by exploring the physical relationship between the endosymbiont intracellular population and this organelle at the cellular level as well as the functional consequences of a *Wolbachia* infection on the ER. To avoid cell line-specific phenotypes and to take in account the impact of the host genetic background, two cell lines showing different gene expression profiles have been infected with the same wMel strain. Specifically, live studies and observation of fixed cells reveal a complex and dynamic interaction between wMel and the ER. This organelle, and not the Golgi apparatus as previously suggested, appears to be the source of the endosymbiont vacuolar membrane. *Wolbachia* redistribute the ER without triggering pathological morphologies. In addition, gene expression analyses indicate that UPR and ERAD key players are not upregulated upon *Wolbachia* infection, and immunostaining studies of ubiquitin chains with degradative roles confirm that ERAD-derived proteasomal degradation is not increased, suggesting that *Wolbachia* do not induce ER stress and proceed through subversion of other host pathways to salvage amino acids.

## Results

### The host genetic background influences the *Wolbachia* titer in *Drosophila* cell cultures

In order to gain insights into the general mechanisms of host cell subversion operated by the *Wolbachia* strain wMel in its natural host *D. melanogaster* to sustain its intracellular lifestyle, and to minimize cell line-specific phenotypes, we established new wMel infections in two *D. melanogaster* cell lines described to display distinct gene expression profiles [24]. The two selected cell lines are adherent, facilitating cellular analyses on live and fixed samples. While both cell lines express about 6,000 genes, nearly half of them show considerable expression variations between cell lines. 1182-4H is an acentriolar haploid cell line derived from maternal haploid *mh 1182* mutant embryos [25,26]. S2R+ are tetraploid male cells derived from the original Schneider’s cell line [27,28]. We chose to introduce in these two different genetic backgrounds a wMel strain derived from JW18, very closely related to the wMel genome of reference [28]. The infected JW18 cell line has been commonly used in numerous studies as a reference cell line to explore the *Wb-host* interactions and the *Wb-* induced viral protection at the cellular and molecular levels [19,28–31]. To infect naive cell lines, wMel bacteria were purified from JW18 cell cultures and added to flasks of uninfected 1182-4 and S2R+ cells (See Methods). JW18 cells harbor fluorescent GFP-Jupiter decorated microtubules. This helped us to confirm the exclusion of cell contaminant during the infection process. After one month, we found the infection to be partial in both cell lines, and an infection dynamics time course experiment confirmed the slow progress of the infection (S1A Fig). Another round of infection was then repeated, leading to stably infected cell lines as determined by immunofluorescence with an anti*-Wolbachia* surface protein-WSP-antibody (See Methods and Fig 1A to C’), named hereafter 1182-4 *Wb* and S2R+ Wb. The vast majority of cells is infected in 1182-4 Wb, and the infection is total in S2R+ Wb. The *Wb* titer is also much higher in S2R+ *Wb* compared to 1182-4 *Wb,* reaching several hundreds of endosymbionts per individual cells (Fig 1B’,C’; S1 and S2 movies). These high *Wb* titers do not significantly affect the host cell viability (Sup1B Fig). We used the WSP-associated fluorescence area, expressed as a percentage of the total cell surface, acquired from full confocal image projections as a proxy to quantify the *Wb* titer in both cell lines (Fig 1D). We concluded that the S2R+ genetic background is more permissive to the wMel infection.

**Fig.1.**
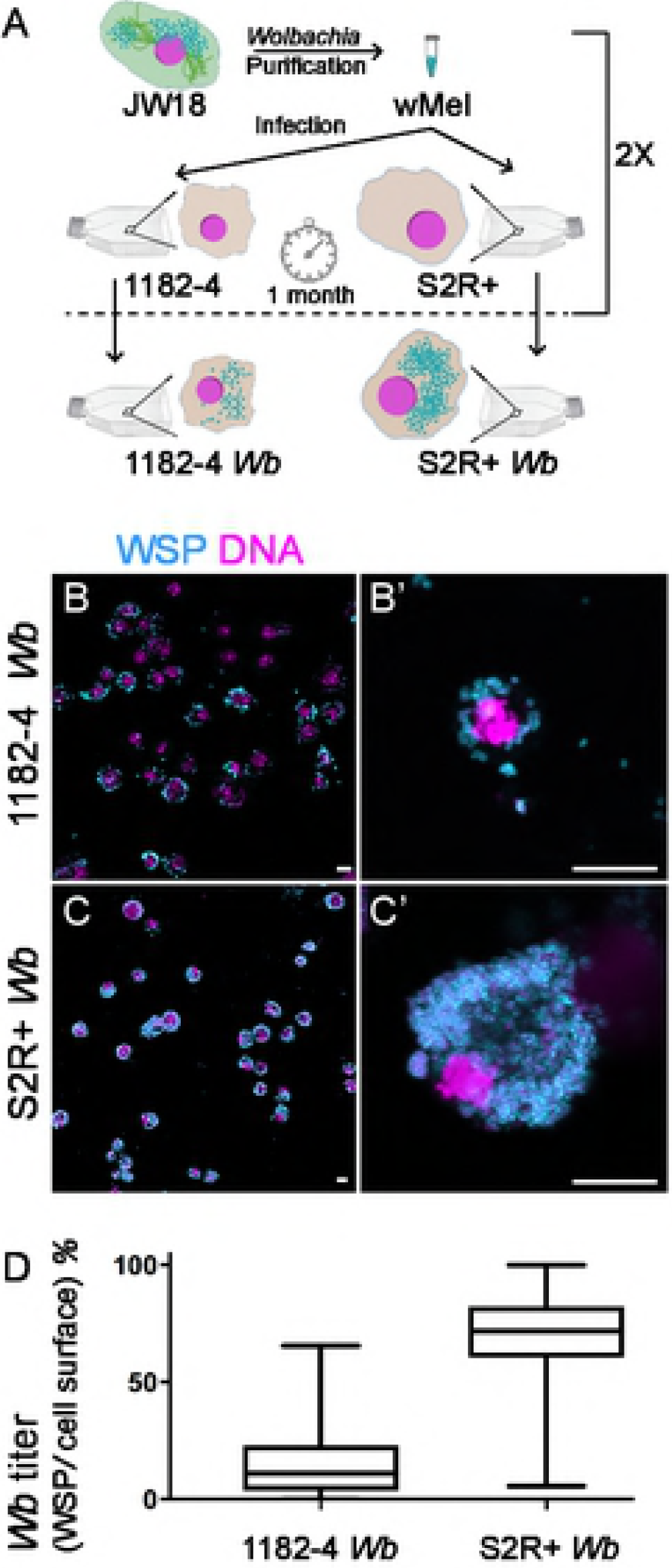
Influence of the genetic background of *Drosophila* cell lines on *Wolbachia* wMel titers. (A) Summary of the experimental approach to infect 1182-4 and S2R+ cell lines. See Materials and Methods. (B to C’) Confocal acquisitions of infected cell lines immunostained with an anti-WSP decorating the *Wolbachia* surface in cyan, DAPI is in magenta. (B’) and (C’) are higher magnifications showing individual cells corresponding to Supplemental movies 1 and 2. Scale bar = 10 microns. (D) Box plot graphs of *wMel* normalized titers in 1182-4 *Wb* and S2R+ Wb, expressed as a percentage of fluorescence surface associated with the anti-WSP staining per cell surface area (n=181 and median= 11% for 1182-4 *Wb,* and n=215 and median= 71% for S2R+ Wb).

### The Golgi apparatus distribution and morphology are not affected by the presence of *Wolbachia*

Using a moderate and variable *Wb* titer in 1182-4 *Wb* on one hand, and a remarkably high *Wb* titer in S2R+ *Wb* on the other hand, we sought to describe the influence of the *Wb* endosymbionts on the host cell physiology, taking into account the *Wb* level. The subcellular distribution of organelles is tightly linked to their function [32], and can be affected together with their morphology, by intracellular pathogens [33]. The *Wb* reside into a vacuole made of a host-derived membrane. Previously *Wb* and the Golgi cisternae were described to reside in the same subcellular compartment close to centrioles in the *Drosophila* embryo, therefore the Golgi apparatus has been proposed to be the source of the Wb-containing vacuole [34]. Moreover the Golgi apparatus can be subverted and fragmented by intracellular pathogens such as *Chlamydia,* that are surrounded by Golgi ministacks to facilitate lipid acquisition [35]. We reasoned that the amount, the localization and the morphology of the organelle providing membranes to the Wb-containing vacuoles may be potentially affected in a *Wb* titer-dependent manner. To investigate the relationship between *Wb* and the Golgi apparatus, S2R+ and acentriolar 1182-4 cells were both co-stained with an anti-*Wb* surface protein-anti-WSP-and a cis-Golgi marker-GM130-, in presence and absence of endosymbionts (Fig 2A). The Golgi apparatus typically displays cell type-specific patterns, and the cis-Golgi often appears as large foci in 1182-4 cells, and as many smaller foci in S2R+ cells. When the *Wb* do not fill the entire cytoplasm, i.e. in 1182-4 *Wb* cells, a thorough visual inspection did not allow us to draw a correlation of subcellular localization between the endosymbionts and the Golgi apparatus. In addition, the number and size of GM130-positive foci did not appear influenced by the abundance of *Wb* endosymbionts in either infected cell lines (i.e. Fig 2A dashed lines for cells with either high or low *Wb* levels, and B). Unlike in a previous report establishing the Golgi apparatus as a source of vacuolar membrane, we never observed GM130-positive *Wb* vacuoles [34]. We next checked the morphology of the Golgi apparatus in presence of *Wb* by ultrastructural studies (Fig 2C). The Golgi cisternae appeared properly arranged, and we could not detect any morphologies that would differ from non-infected cells, despite heavy loads of endosymbionts in the S2R+ *Wb* cell line. Together, this data set suggests that the Golgi apparatus does not appear to be subverted by *Wolbachia* at the subcellular level, and does not support the hypothesis of this organelle being a source of membrane for the endosymbionts.

**Fig.2.**
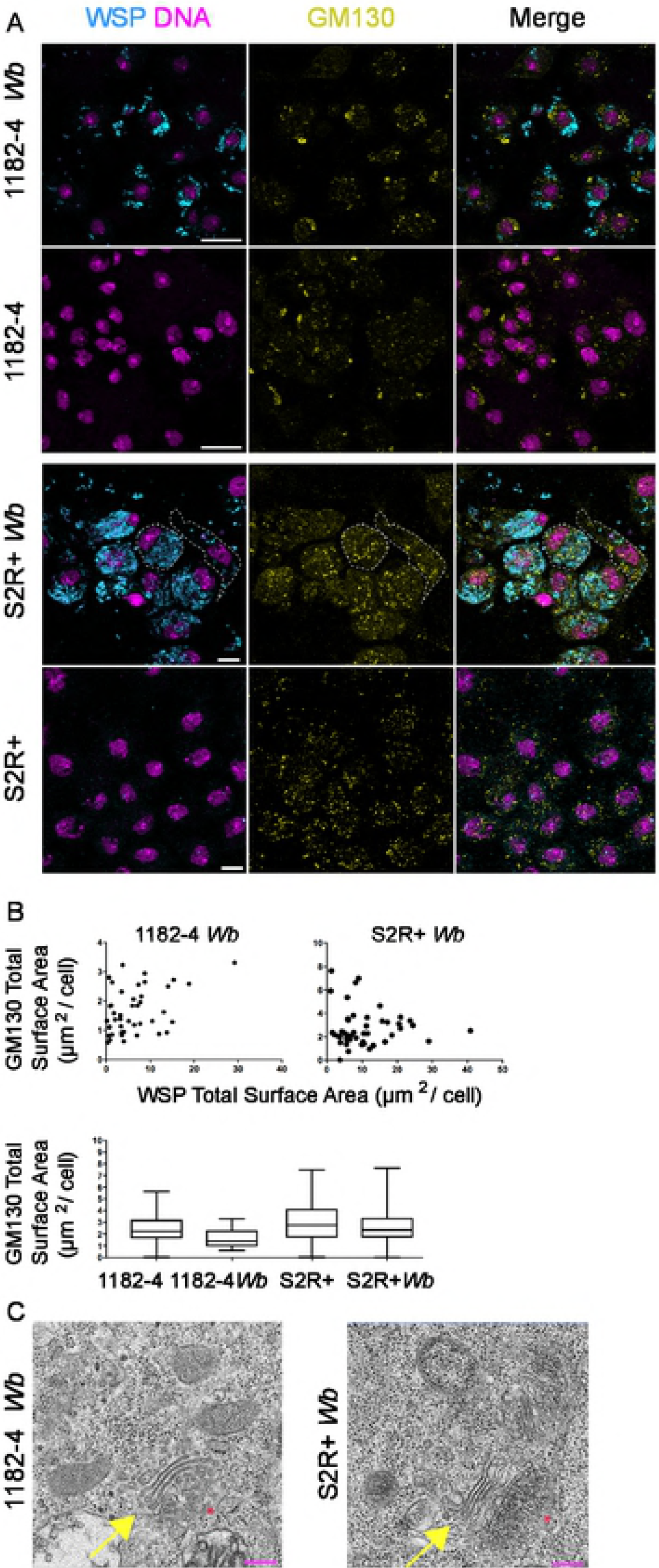
*Wolbachia* subcellular localization and titer do not influence the Golgi apparatus distribution and morphology. (A) Confocal acquisitions of infected cell lines immunostained with an anti-WSP-decorating the *Wolbachia* surface in cyan-, with GM130-yellow-. DAPI is in magenta. Scale bars= 10 microns. Dashed lines encompass in S2R+ *Wb* a highly infected cell-left cell-and a infected cell at low level-right cell-(B) Top graphs: Distribution of GM-130 foci sizes in function of the *Wb* density measured on full projections of confocal images (n= 44 cells for 1182-4 *Wb* and n=46 cells for S2R+ Wb). Bottom graph: Amount of cis-Golgi expressed as GM130 total signal per cell measured on full projections of confocal images, in infected and noninfected cell lines (n= 1250 cells for 1182-4; 797 for 1182-4 *Wb,* and n=743 cells for S2R+ cells and n=864 for S2R+ Wb). (C) Transmission electron microscopy images of the Golgi apparatus in *Wolbachia-infected* cells. The Golgi stacks-yellow arrows-appear normal (n>10, the red stars indicate the trans-Golgi). Scale bars= 200 nm

### *Wolbachia* interact with the Endoplasmic Reticulum, source of their vacuolar membranes

A previous study based on electronic microscopy has reported observations of *Wb* in close contact with ER tubules, and in some instances a continuum between the ER and the *Wb* vacuolar membrane [19]. To better understand how and to what extent the *Wb* intracellular population interact globally with the ER, we performed simultaneous live observations of the endosymbionts and of this organelle. To this end, we used the SYTO 11 DNA live dye that stains preferentially *Wb* [36], and an ER tracker, that recognizes the sulfonylurea receptors of ATP-sensitive K+ channels located on ER membranes. We first performed confocal time lapse fluorescence imaging of 1182-4 *Wb* cells. Cortical areas enriched in tubular ER were chosen for time lapse analyses because they offer a better resolution of these dynamic structures (Fig 3A). We typically observed three categories of Wb. Some peripheral *Wb* clusters did not show any obvious interactions with the ER (Fig 3A grey arrows), some were juxtaposed to the ER and displayed tightly coupled dynamics (Fig 3A orange arrow), while few *Wb* appeared to be localized within dynamic ER tubules (Fig 3A yellow arrowhead, and see S3 movie that recapitulates these observations). We next used the same fluorescent markers in 1182-4 *Wb* and S2R+ *Wb* cells to score the different types of interaction between *Wb* and the ER (Fig 3B). Striking differences appeared in these two different cellular environments. While in random focal planes 62% of *Wb* did not reside in close ER vicinity in 1182-4 *Wb* cells, only 2% were distant from the ER in S2R+ *Wb* cells. Hence a majority-80%-of endosymbionts were in close contact with the ER in S2R+ Wb, while only 34% contacted the ER in the 1182-4 genetic background. Interestingly 17% in S2R+ *Wb* and 9% in 1182-4 *Wb* appeared either inside the ER and/or surrounded by an ER tracker-positive membrane (Fig 3C). Together this dataset shows that the physical interaction of *Wolbachia* with the ER is highly dynamic. The presence of ER tracker around some endosymbionts strongly suggests that this organelle is a source of vacuolar membrane. Some *Wb* were detected in ER tubules, and only a minority of endosymbionts display an ER tracker-positive vacuolar membrane, leading us to hypothesize that they may represent newly acquired membranes, whose composition is subsequently modified by *Wb* (i.e. less or no ATP-sensitive K+ channels leading to ER tracker-negative *Wb* vacuoles). In addition, time lapse recordings showing Wb-ER coupled dynamics reveal a tight physical interaction between the *Wb* vacuole and this organelle, suggesting potential signaling events and/or possible nutrient uptake. The increased association of *Wb* with the ER in a S2R+ genetic background, highly permissive to the *Wb* infection, suggests that the ability to subvert the ER is crucial for *Wolbachia* to thrive intracellularly.

**Fig.3.**
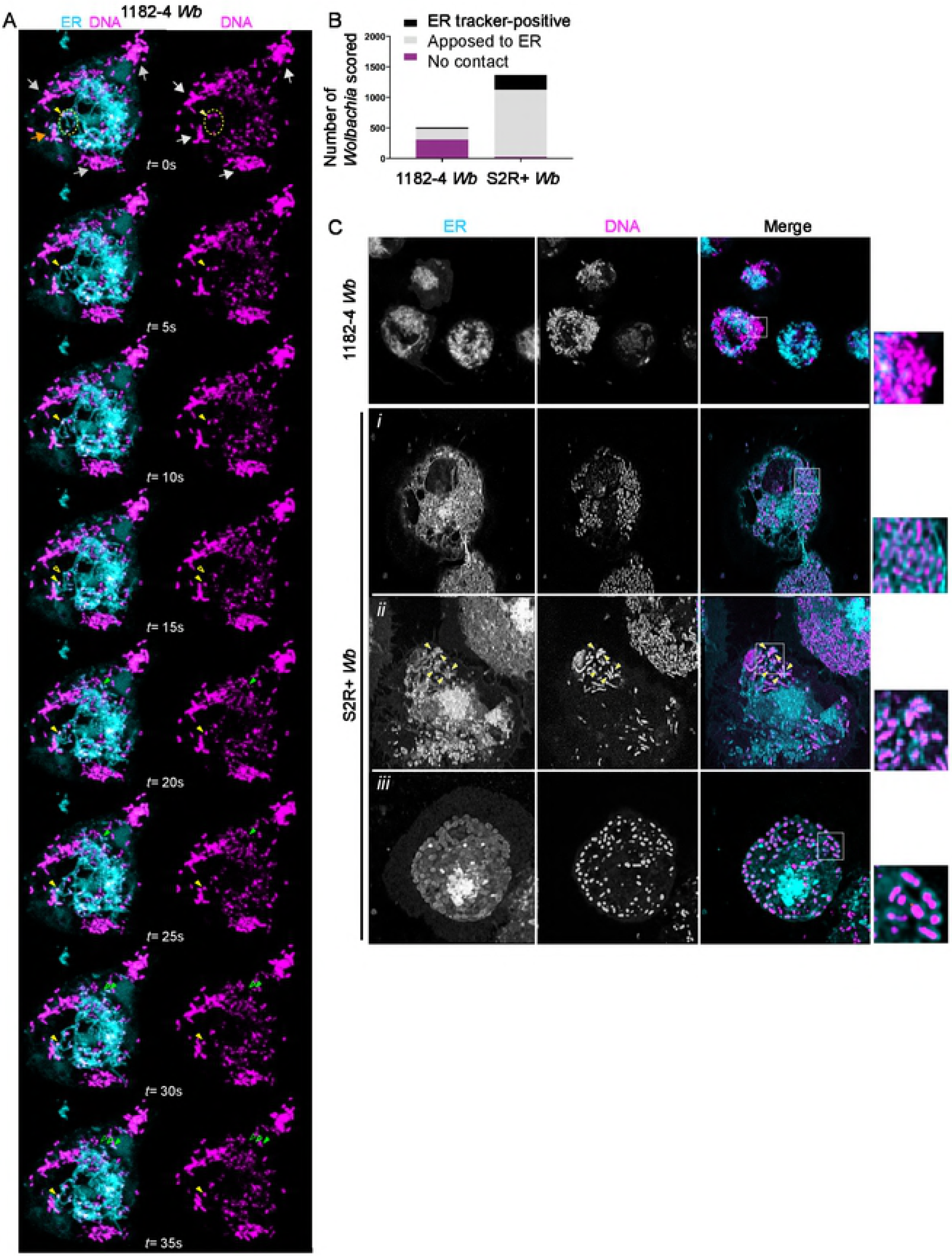
*Wolbachia* physically interact with the endoplasmic reticulum. (A) Time-lapse acquisitions at a surface focal plane in a 1182-4 *Wb* cell stained with the DNA dye SYTO 11-magenta-to highlight Wb, and an ER tracker-cyan-. A t=0 second, grey arrows point to peripheral *Wb* clusters that are not in close contact with the ER. The orange arrow points towards some *Wb* remaining in close contact with the ER during the time lapse duration. The dotted yellow circle highlights some *Wb* located within ER tubules. A single *Wb* within an ER tubule is tracked by the plain yellow arrowhead, and its previous position is indicated by an empty yellow arrowhead (i.e. at t=15s). Similarly, the movement of a single *Wb* surrounded by an ER-derived membrane is tracked by green arrowheads (t=15s to t=35s). See the corresponding supplemental movie 3. (B) Scoring of *Wb-* ER interactions, observed with SYTO 11 and the ER tracker in 1182-Wb and S2R+ *Wb* cells, in random focal planes of n= 18 and n= 12 cells respectively. Bacteria co-localized with ER tubules, or surrounded by an ER tracker-positive membrane were counted as “ER-tracker positive”. (C) The different interactions between *Wb* and the ER are highlighted on these confocal images, with clusters of *Wb* not in contact in 1182-4 *Wb*-see inset-. The following rows are different examples in S2R+ cells showing i) *Wb* in close contact with the ER, ii) a *Wb* cluster composed of individual *Wb* surrounded with an ER tracker-positive membrane-yellow arrowheads-; iii) and in rare instances all individual *Wb* of the cell being surrounded with an ER tracker-positive membrane.

### The ER subcellular distribution is affected by *Wolbachia*

Because the ER*-Wb* contacts are prominent in S2R+, we first examined the ER by confocal microscopy to assess the impact of *Wb* on its distribution. In non-infected cells, the ER appears principally composed of a dense perinuclear network of tubules and vesicles, while cisternae are less detectable. The cell periphery and cortical areas are enriched with ER tubules, which are often branched (Fig 4A). In contrast, in infected cells a fraction of the ER becomes heavily clustered close to the nucleus (Fig 4A cyan arrows), while cytoplasmic regions harboring *Wb* are highly enriched in tubular ER (Fig 4A yellow arrowheads and bottom row).

**Fig.4.**
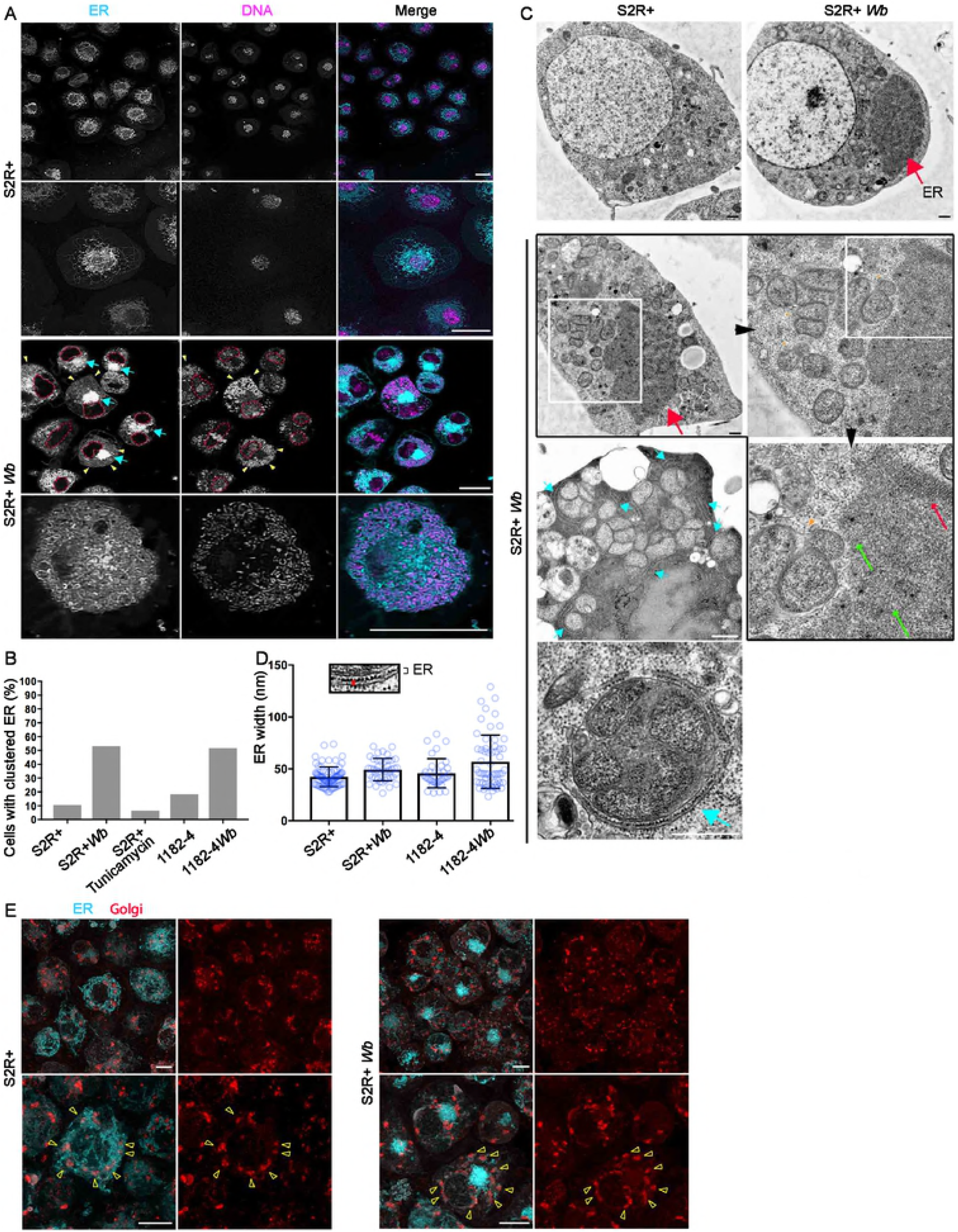
*Wolbachia* impact the ER distribution but not its structure in both S2R+ *Wb* and 1182-4 *Wb* cells. (A) Live imaging of S2R+-top rows-, and S2R+ *Wb*-bottom rows-stained with SYTO 11-magenta-and the ER tracker-cyan-. For S2R+ *Wb* cells, dashed lines encompass the nuclei, yellow arrowheads the colocalization of *Wb* and ER tubules, and blue arrows point to the clustered ER. The last row is a cortical focal plane showing the intense ER tubular network associated with Wb. Scale bar= 10p.m. (B) Occurrence of clustered ER in various cell lines, with the addition of the S2R+ cell line treated with Tunicamycin at 10pg/mL for 48 hours. For S2R+ n= 179; S2R+ *Wb* n= 123; S2R+ with Tunicamycin n=227; 1182-4 n=76; 1182-4 *Wb* n=120. (C) Electron micrograph of S2R+ and S2R+ Wb. The top row highlights the presence of a darker ER mass-red arrow-, numerous *Wb* are visible in between the nucleus and the ER cluster. The second row is a series of consecutive enlargements of an ER cluster in the vicinity of vacuoles containing multiple *Wb*-orange arrowheads pointing to the vacuolar membrane-. Green arrows and the red arrow indicate the tubular ER and piled ER membranes respectively. Cyan arrows point towards ER membranes encompassing vacuoles containing multiple *Wb* in the cell periphery. The last image depicts a single vacuole with multiple *Wb,* tightly surrounded by rough ER-cyan arrow-. Scale bar= 500 nm. (D) The ER inter-membrane distance in the different cell lines. Measurements were taken on high magnification electron micrographs as depicted-red line-, and the average thickness varies from 42 to 56 nm. For S2R+ n=75; S2R+ *Wb* n=43; 1182-4 n=35; 1182-4 *Wb* n=58. (E) Live imaging of S2R+ and S2R+ *Wb* cells stained simultaneously with ER-cyan-and Golgi-red-fluorescent trackers. Upper panels are lower magnifications and lower panels are higher magnifications. Arrowheads point towards Golgi foci. Scale bar=10pm.

We defined this mass of ER as “ER clusters”, which is greatly enhanced by the presence of *Wb* in both cell lines (Fig 4B). We wondered whether this ER distribution was a consequence of an ER stress, and S2R+ were treated with tunicamycin, an ER stress inducer, which did not increase the occurrence of this phenotype compared to untreated S2R+ cells (Fig 4B). ER morphological aberrations that may not be detectable by confocal fluorescence microscopy have been reported in *Wb*-infected cells such as ER tubule swelling and an increase in cisternae [19], leading us to perform EM ultrastructural studies on s2R+ *Wb* and 1182-4 *Wb* cells, and on their naive counterparts (Fig 4C,D). The dark ER mass is easily distinguishable in infected cells-thick red arrows-. A closer look at this cluster reveals it is composed of randomly-thin green arrows-and orderly-thin red arrow-packed tubules or cisternae. No swollen structure was detected within these clusters in either cell types. In the periphery, multiple *Wb* share very often a same vacuole, tightly apposed to rough ER-cyan arrows, and bottom image-. Incidentally, these multi-*Wb* vacuoles were encountered much more frequently in the highly permissive S2R+ genetic background compared to 1182-4. We then searched for a size increase of cisternae and swollen ER tubules without success in S2R+ *Wb.* Measurements of ER inter membrane distances by electron microscopy however revealed very marginal ER swelling in 1182-4 Wb, not affecting the average thickness of ER in this cell line (Fig 4D). Last, because of the dramatic ER redistribution observed in S2R+ *Wb* occurring in more than half of these infected cells, we investigated at the individual cell level the impact of this ER defect on the Golgi apparatus distribution (Fig 4E). In non-infected cells, the Golgi foci are surrounded throughout the cell periphery by large amounts of the ER (Fig 4E left upper and lower panels, yellow arrowheads point to Golgi foci). In Wb-infecting cells showing ER clusters, the Golgi units do not coalesce toward the ER mass (Fig 4E right panel top images), and their distribution is not appear significantly perturbed. Although they remain associated with some ER (Fig 4E yellow arrowheads on bottom images), the overall distance between most of the ER and the Golgi apparatus is increased. In conclusion, *Wolbachia* dramatically redistribute the ER without affecting its luminal width, since we did not observe any ultrastructural variations in presence of the endosymbionts. The high titer in S2R+ *Wb* correlates with a tight association of *Wb* with the ER, and in general a large fraction of this organelle becomes spatially restricted, close to the nucleus, upon a *Wb* infection. This defect could potentially affect its function and interactions with other organelles such as the Golgi apparatus. Attempts to phenocopy the ER compaction with tunicamycin did not succeed, suggesting that this redistribution is operated by *Wb* independently of a potential ER stress.

### *Wolbachia* do not induce ER stress in 1182-4 and S2R+ genetic backgrounds

We next sought to examine the impact of a *Wb* infection on the ER functions. To ensure protein homeostasis in the cell, one of the role of the ER is to control the proper folding and maturation of proteins through the unfolded protein response-UPR-, upregulated when misfolded proteins accumulate. When these adaptive responses are not sufficient, the endoplasmic-reticulum-associated protein degradation-ERAD-pathway is in turn activated to target and retrotranslocate ER misfolded proteins to the cytosol, where they are addressed towards a degradation pathway by the ubiquitin-proteasome machinery [37].

We first checked whether the ERAD function was subverted in order to provision *Wb* with amino acids derived from an increased proteolytic activity, as previously suggested in Wb-infected JW18 cells [19]. We first stained cells with the FK2 antibody recognizing all mono-and polyubiquitylated proteins, but not the free ubiquitin, considered as a good proxy to assess proteasomal degradation-associated polyubiquitylation marks-K48 and K11 poly-Ub-, since these degradation marks are the most abundant among polyubiquitylated chains in the cell [38] (See Methods and Fig 5A). We quantified the total fluorescence surface associated with the polyubiquitylation foci on full confocal projections, and we found the presence of *Wb* to correlate with 2.5 and 4.2 times as many polyubiquitylation in 1182-4 and S2R+ genetic backgrounds respectively (Fig 5A,B). We reasoned that a proteasomal degradation-linked poly-Ub signal, reflecting a Wb-dependent amino acid demand, should vary according to the endosymbiont titer, that is variable between cells in a given infected cell line. We chose the 1182-4 *Wb* cell line showing fewer heavily infected cells to perform a linear regression highlighting the amount of FK2 foci in function of an increasing *Wb* titer (Fig 5C). We found no correlation between the *Wb* titer and the number of FK2 foci. This suggests that the observed FK2 signal is unlikely to account for an increased proteasomal degradation. To verify this result, we next checked specifically the levels of K11 poly-Ub chains by immunostaining analyses. K11 is the ubiquitin linkage primarily generated by the ERAD pathway [39]. We failed to detect any differences between infected and non-infected cells (Fig 5D). In the fraction of S2R+ *Wb* cells endowed with high *Wb* levels, the ER becomes clustered in an area from which the endosymbionts are excluded. Focusing our attention on these areas to detect a possible enrichment of ER-associated K11 poly-Ub, we did not detect an increase of this ERAD-associated degradation mark (Fig 5D, dashed yellow circle). Both K11-and K48-linked poly-Ub chains are involved in ERAD [40], therefore we checked the levels of K48 poly-Ub, that also appeared undistinguishable in infected cells compared to their non-infected counterparts (S2 Fig.). Together these results indicate that the global increase of cellular polyubiquitylation in presence of *Wolbachia* does not reflect an increase in proteasomal degradation-associated K11/K48 polyubiquitylation marks.

**Fig.5.**
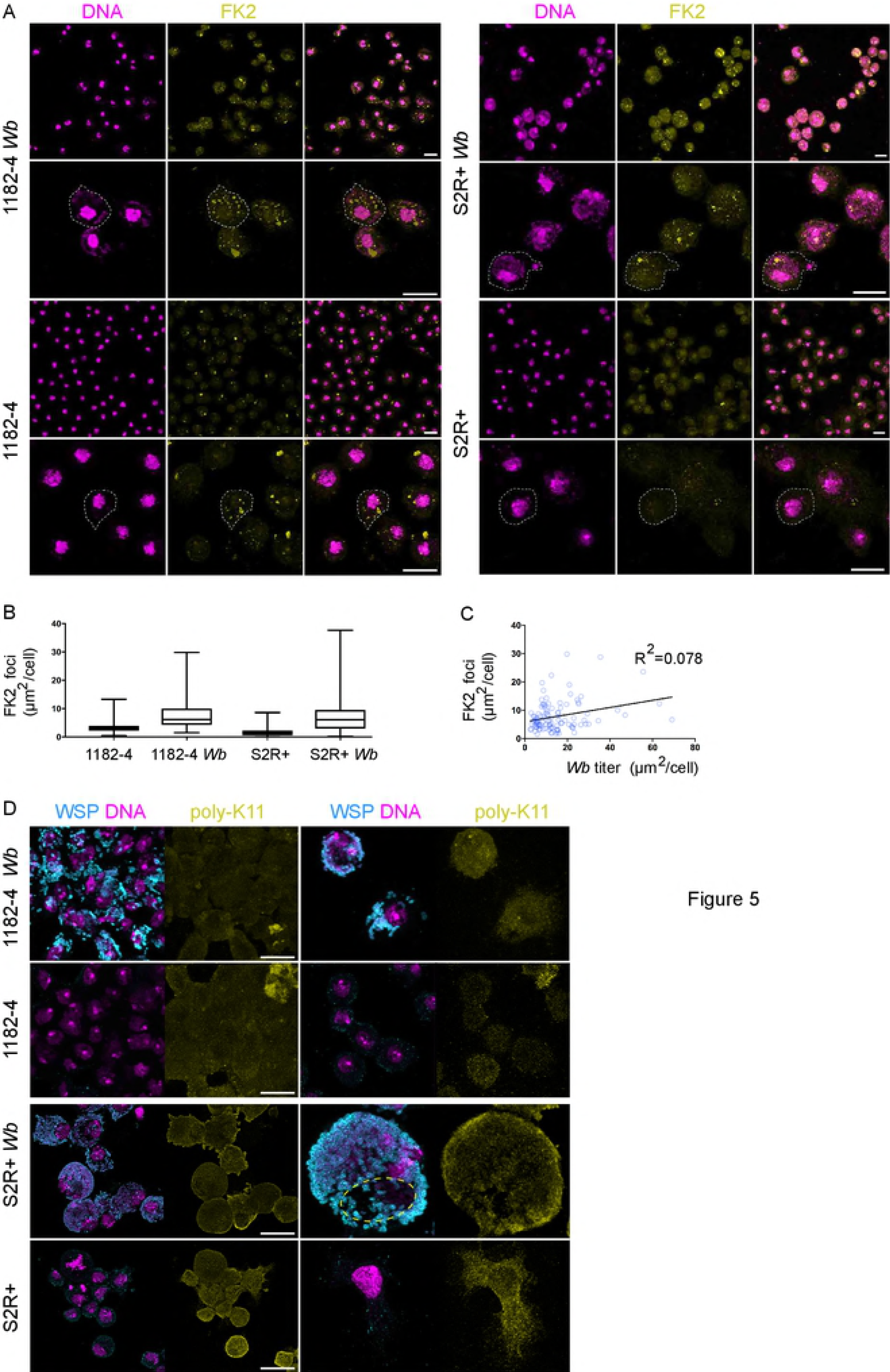
Polyubiquitin linkages associated with ERAD and proteosomal degradation are not increased in presence of *Wolbachia.* (A) Confocal acquisitions of the infected and non-infected 1182-4 and S2R+ cell lines stained with DAPI-magenta-and the monoclonal antibody FK2-yellow-, recognizing all mono-and poly-ubiquitylated proteins, but not free ubiquitin. Dashed lines encompass individual cells, scale bar=10 pm. (B) Box plot graphs showing the FK2-positive foci quantification, expressed as total areas per full confocal projections in individual cells. For 1182-4 n=218; 1182-4 *Wb* n=106; S2R+ n= 212; S2R+ *Wb* n= 195 cells. (C) Linear regression of the FK2-positive total area per 1182*-Wb* cell, in function of the *Wb* titer, established on the DAPI signal (cf. Materials and Methods), n=104 cells. (D) Confocal acquisitions of the infected and non-infected 1182-4 and S2R+ cell lines stained with WSP-magenta-and an anti-K11-linkage polyubiquitin-yellow-. The dashed line highlights the cell area of a heavily Wb-infected S2R+ *Wb* cell, containing a mass of ER, physically excluding the endosymbionts. Scale bar= 10p.m.

We decided to perform quantitative PCR analyses to investigate the UPR and ERAD responses at the gene expression level in the presence of Wb, in order to characterize the level of ER stress potentially generated by the endosymbionts. Briefly, upon a stress leading to accumulation of misfolded proteins, the ER transmembrane stress sensors PERK, ATF6, and IRE1 release the chaperone Bip in the ER lumen, and an UPR response is activated. This response aims at decreasing protein translation and enhancing the protein folding capacity in the ER, by upregulating the expression of chaperones and UPR sensors (Fig 6A and [41]), while the ERAD pathway drives misfolded protein to undergo proteolysis. We first selected *D. melanogaster* genes confirmed to respond to tunicamycin-induced ER stress, and that are involved in both UPR and ERAD responses [42]. We next monitored these candidate genes in the 1182-4 genetic background by submitting the cell line to a tunicamycin treatment for 48 hours at 10 [μg/mL (Fig 6B). We found a ~2 fold gene expression upregulation for the three UPR sensors *perk/gcn2, atf6* and *ire1* (Fig 6A, B top graphs). In addition, a number of ERAD key players, the *derlin* orthologs *der-1* and *der-2, sel1L/hrd3 and hrd1/sip3* whose products associate to form a complex, as well as members of the ubiquitin ligase complex were upregulated from 3 to more than 5 folds. With this experiment validating the 1182-4 cell line responsiveness to ER stress, we next measured the impact of *Wb* on this stress in the 1182-4 *Wb* (Fig 6B bottom graphs). We did not detect any induction of the UPR sensors or downstream targets. Similarly, none of the ERAD key players that responded to tunicamycin were affected by the presence of Wb. This shows that *Wolbachia* do not trigger an ER stress response leading to increased UPR and ERAD activities in 1182-4 *Wb* cells. Last, we verified the level of ER stress in S2R+ *Wb* cells using a fluorescent ATF-4 activity reporter gene-the translational inhibitor 4E-BP-that responds to the PERK/GCN2-ATF4 pathway through ATF4 binding sites [43]. The fluorescence was monitored 48 hours after transfection with the 4E-BP intron dsRed reporter, and a tunicamycin treatment was added as a positive control of ER stress (Fig 6C, and Methods). Transfected cells showed in presence of tunicamycin high nuclear and cytoplasmic fluorescence levels. Quantification of the fluorescence revealed a level of ATF4 signaling activity upon ER stress 4 times higher on average compared to non-treated S2R+ cells. The fluorescence levels expressed in S2R+ *Wb* cells appeared similar to what observed in S2R+ cells, suggesting that the presence of *Wb* do not cause a significant stress in the S2R+ genetic background. Altogether, this data set suggests that in these two host cell genetic backgrounds, the *Wolbachia* can proliferate and persist in a stable manner without triggering ER stress and in particular the ERAD pathway, implying that other mechanisms than ERAD-induced proteolysis should exist to salvage amino acids.

**Fig.6.**
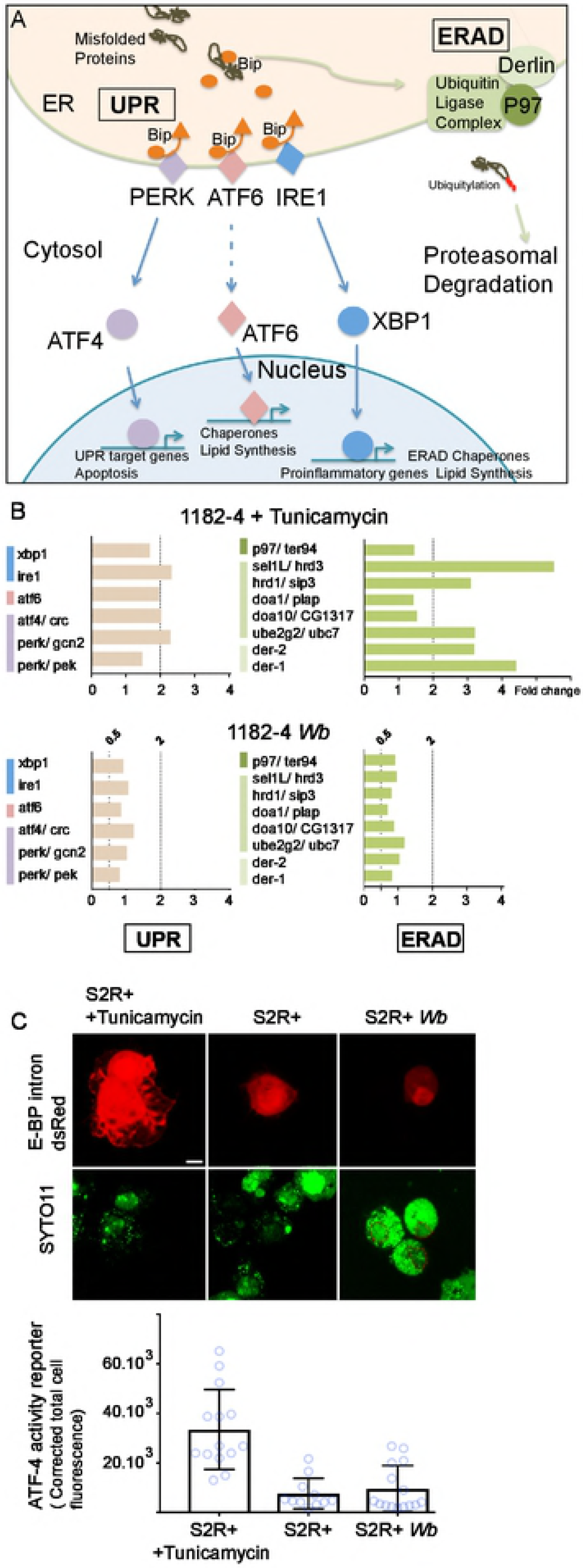
*Wolbachia* do not induce induce ER stress. (A) Schematic summary of the UPR and ERAD pathways. The color code highlights the three UPR pathways and the ERAD and is identical to what employed in (B). (B) Genes tested by quantitative PCR, in presence of tunicamycin-top graphs-, or Wolbachia-bottom graphs-. UPR genes are on the left and ERAD genes on the right. Gene expression fold changes are represented, and variations comprised between a 2-fold increased expression-”2” above the dashed line-and a 2-fold decreased expression-”0.5” are considered insignificant. (C) S2R+ cells transfected with the ATF4 activity reporter E-BP intron dsRed; after a 48hr-long treatment with tunicamycin at 10 [ig/mL, or in presence of *Wb.* DNA is stained with SYTO-11-green-. In absence of *Wb,* nuclei incorporate the dye at various levels, while in presence of Wb, the dye stains preferentially the endosymbionts compared to the nuclei, highlighted with a red dashed line. Two adjacent transfected cells are shown in presence of tunicamycin, and only one for S2R+ and S2R+ Wb. The graph represents quantifications of the dsRed fluorescence levels in each conditions. For S2R+ n=14; S2R+ with tunicamycin n=11; and for S2R+ *Wb* n=15.

## Discussion

A number of studies these past years have started to investigate the basis of the *Wb* intracellular lifestyle and their impact on the cell homeostasis using *in vitro* cell culture models (i.e. [13,16,18,19,44,45]). The results of these studies can be variable depending on the *Wb* strain and the infected insect cell lines. In order to minimize the bias of a cellular context potentially leading to cell line-specific phenotypes, we infected two genetic backgrounds presenting an important variation at the level of the expressed genes [24]. Additionally, the two cell lines were infected with a single wMel strain, that derives naturally from *D. melanogaster.*

Here we identified the endoplasmic reticulum as a source of vacuolar membranes for *Wolbachia* in *D. melanogaster* species, and we observed close appositions between the replicative vacuole of these endosymbionts and the ER membrane. These appositions are likely to lead to the biogenesis of ER-derived *Wb* vacuoles, while sometimes allowing fusion with this organelle. Coupled dynamics between *Wb* and the ER tubules seen in time lapse microscopy reveals tight and prolonged interactions, supporting as well the possibility of nutrient uptake from the ER. The cellular context greatly influences the *Wb* titer, and a permissive environment correlates with more apposition events with the ER, suggesting that the ability of *Wb* to subvert the ER in a given environment correlates with growth and replication. A *Wb* infection redistributes the ER, and while a tubular network associates with the endosymbionts, a significant fraction of this organelle shrinks to become compacted close the cell nucleus. Although the functional impact of this ER clustering remains unclear, the ultrastructural ER organization does not reveal swollen compartments or more cisternae. Gene expression analyses of central ER stress players, as well as immunofluorescent studies of ERAD-induced proteolysis key marks indicate that the Wb-induced ER subversion does not trigger the UPR nor an increased proteolysis. Hence the *Wb* level, whether low or high, does not seem to perturb the ER-regulated mechanisms of cell homeostasis in a significant manner. Incidentally, these results indicate that *Wb* is likely to rely on other sources than ERAD-induced proteolysis to salvage amino acids.

The *Wolbachia* endosymbionts are transmitted vertically in their arthropod or filarial nematode hosts, from mothers to their offspring. Once in the egg they next colonize specific somatic tissues and the germline during embryonic and larval developmental stages, following asymmetric segregation during cell mitotic divisions [2]. Although a germline tropism has been described, implying that *Wb* can pass from cell to cell either artificially in *Drosophila* through abdominal injections of purified Wb, or through a developmentally regulated colonization of the filarial nematode ovary [46,47], they do not share with most intracellular pathogens the ability to easily infect naíve cells, thus limiting their horizontal transfers. It has been demonstrated that *Wb* can pass from infected to non-infected cell in *in vitro* assays, without requiring cell-to-cell contact, possibly through secretion [30]. If active mechanisms of cell entry are not precisely described, passive uptake mechanisms through phagocytosis explain at least in part their entry in cell culture assays. To optimize the infection of naive cell lines, we set up a protocol of *Wb* enrichment from a Wb-infected cell culture. This allowed us to expose cells to very high bacterial concentrations. Although *D. melanogaster* cell cultures have a strong capacity of engulfment–which does not make them an ideal model to study mechanisms of bacterial cell entry-, artificial infections of naiíve cell culture with *Wolbachia* remain nonetheless a slow process. The fact that a significant proportion of cells remained uninfected after one month suggests indeed that extracellular *Wb* originating from possible secretion or dead cells do not have strong infection capacities and that colonization of a naíve environment remains a challenge. This is in part due to their slow replication cycle estimated to last 14 hours [48], but it is also very likely that some *Wb* do not succeed in escaping autophagy. Those nonetheless succeeding at surviving and replicating not only need to modify the phagosome membrane along the endocytic pathway to avoid the cell surveillance, but also need to acquire new membranes and nutrients.

The ER represents a nutrient-rich compartment devoid of antimicrobial functions, and several intracellular bacteria derive their vacuole from, and/or replicate in, this organelle [49]. Such is the case of *Legionella pneumophila* and *Brucella abortus* that possess like *Wb* a type IV secretion system they employ upon infection to secrete an array of effectors subverting cellular machineries to gain access to ER. *L. pneumophila* regulate membrane trafficking through modulations of GTPase signalling pathways interfering with early secretory vesicles to ultimately allow fusion of the *Legionella* vacuole with ER-derived membranes [50]. Along the endocytic pathway, *B. abortus* co-opt the ER exit sites-ERES-, involved in the vesicular trafficking towards the Golgi., thus acquiring an ER-derived vacuolar membrane [51]. Similar to observations of these pathogens, our ultrastructural studies have revealed a tight association of *Wb* with rough ER membranes. In addition, live experiments have demonstrated that some *Wb-* containing vacuoles appear positive for a fluorescent and specific ER tracker, and in some instances *Wb* were located within ER tubules, strongly suggesting that the ER is a source of membrane for Wb. We hypothesize that the presence of ER tracker-negative Wb-containing vacuoles indicates a maturation process in the biogenesis of the membrane surrounding Wb, although we cannot rule out other sources of membranes. The compaction of ER observed in both 1182-4 *Wb* and S2R+ *Wb* cell lines places *Wb* in between ER and the Golgi apparatus, which could potentially favors *Wb* interactions with the ERES. *Wb* could benefit from co-opting the COPII vesicles routing towards the Golgi to acquire membranes, lipids and other nutrients. This is in accordance with the discovery that in presence of the pathogenic strain Wmelpop, cholesterol homeostasis is affected [18]. Not only *Wb* likely incorporate cholesterol into their membranes as a substitute for lipopolysaccharide, but also proper ER-to-Golgi vesicular trafficking requires cholesterol [52]. Hence *Wb* may interfere with the anterograde trafficking. In addition, a lipidomic analysis has shown that the wMel affect the sphingolipid metabolism and deplete mosquito cells from ceramide and derived sphingolipids [16]. Ceramides are synthesized in the ER and exported to the Golgi [53]. They play an important role during bacterial infections as part of a pro-apoptotic lipid signalling [54] and sphingolipids regulate autophagosome biogenesis and endocytic trafficking [55], suggesting that a Wb-induced decreased availability of these lipids may prevent xenophagy and/or apoptosis. It is then possible that the interaction of *Wolbachia* with the ER and the derived intracellular vesicular trafficking plays also a central role in immune escape and control of apoptosis. In S2R+ *Wb* cells, the bacterial titer is exceptionally high compared to other infected insect cell lines, and *Wb* often fill the cytoplasm entirely when observed in confocal microscopy with an anti-WSP staining. In this cellular environment unable to efficiently control the *Wb* titer, electron microscopy analyses revealed a high frequency of poly *Wb-* containing vacuoles, possibly resulting from a limited access to new membranes. It is nonetheless interesting to observe that under these conditions the infection is persistent and does not compromise the host cell viability. Since ER tracke-rnegative *Wb* are often observed in the cell periphery, the interaction with ER may be necessary for an active replication.

*Wb* infections are usually characterized by very high intracellular loads of bacteria, usually above a hundred bacteria per cell, similar to other Rickettsiales. Despite the peculiar relationship between *Wb* and the ER, we did not detect an ER stress above levels found in non-infected cells suggesting that a *Wb* infection either does not require this cell response or is able to prevent it. Moreover, prolonged ER stress leads to cell death and seems incompatible with endosymbiosis [56,57]. This conclusion is in addition justified by several lines of evidence. First, although the ER appears redistributed, we did not detect morphological signs of enhanced ER activities linked to ER stress, such as swollen tubules and cisternae, in contrast to a previous study performed with wMel-infected LDW1 cells [19]. Second, we monitored the gene expression levels for the three UPR sensors, downstream targets, and ERAD key players, either by quantitative PCR or by fluorescent assay approaches. We could not find altered gene expressions indicating that a persistent *Wb* infection triggers an ER stress. Last, immunofluorescence studies of polyubiquitin linkages associated with ERAD-driven proteolysis (K11 and K48 polyUb) revealed that these marks are not increased in presence of *Wb.* Since the monoclonal antibody FK2 targets all covalently linked mono-and poly-ubiquitins, it is likely that the increased amount of FK2 foci in presence of *Wb* corresponds to either mono-ubiquitylated proteins; and/or to proteins decorated with polyubiquitin chains on possibly the five other lysine residues of ubiquitin with non-degradative roles, reported to be involved in: K6-mitophagy-, K27-protein secretion and autophagy-, K63-endocytosis, signalling, activation of NF-kappa-B-; K33-kinase modification-, and K29-lysosomal degradation-[38]. It is hence possible that *Wolbachia,* directly or indirectly, influence a number of cellular mechanisms through modulation of polyubiquitylation-dependent signalling events, and this field remains to be explored. Recent proteomic studies provide conflictual evidence regarding *Wb* and the UPR, possibly due to the differences in the *Wb* stains and the host cells employed. The pathogenic strain wMelpop slightly increases (up to 1.36 fold) some UPR-related genes identified by gene ontology analysis [18] while the wStr infection in *Aedes albopictus* cells rather leads to a decrease of proteins involved in ER protein folding [44]. Nonetheless a genome-wide RNAi screen has revealed the importance of UBC6, an ubiquitin-conjugating enzyme part of the ERAD pathway, to sustain the wMel titer [19]. Although we found no evidence for an increased ERAD-induced proteolysis through ubiquitin-targeted proteasomal degradation in presence of *Wb,* this does not rule out the requirement of intact UPR/ERAD response for *Wb* survival. Alternatively, UBC6 may either be involved in a non-ERAD-related function, or since the *Wb* vacuolar membrane appears ER-derived, these endosymbionts may have subverted an ERAD machinery at the level of their own vacuole. The apociplast of apicomplexan parasites is an organelle derived from an algal endosymbiont that has retooled the host ERAD into an apicoplast-localized ERAD-like protein import machinery [58].

The UPR response can be modulated by intracellular pathogens to their advantage, and the three branches–IRE1, PERK, ATF6-can be individually upregulated or inhibited in order to modulate i.e. the host defense through apoptosis or innate immunity response, or to build a replicative niche [59]. Hence, further investigations will be needed to clarify the role of the UPR in a *Wb* infection. However, the absence of an enhanced ERAD-proteasomal degradation pathway suggests that amino acid salvage does rely on mechanisms other than an increased proteolysis. Several studies have shown that the *Wb* infection decreases the global protein translation in the host cell [28,44]. While the mechanisms are still unknown, TORC1 and insulin pathways regulate protein translation based on environmental conditions, and greatly influence the *Wb* titer in *Drosophila* [60]. Future studies will determine whether *Wolbachia* can directly subvert growth signalling pathways to down-regulate translation and therefore increase the pool of free amino acids.

In conclusion, there is no doubt that in an effort to elucidate the mechanisms of intracellular survival employed by *Wolbachia,* the comprehension of subversion strategies will be key: how are ubiquitylation pathways modulated and what are their targets? How do *Wb* acquire ER-derived membranes on one hand, and how do they modulate signalling or synthesis pathways to acquire amino acids and lipids on the other hand? These are the next questions to be addressed. In parallel, the current growing efforts to express the putative *Wb* effectors into surrogate systems, yeast or *Drosophila* cell cultures, should accelerate our knowledge of one of the most commonly encountered endosymbiont.

## Methods

### Cell lines

All the cell lines are derived from primary cultures of *D. melanogaster* cells. JW18 is a kind gift from William Sullivan [61], 1182-4 was obtained from Alain Debec [25,26], and S2R+ from François Juge [27]. JW18, 1182-4, and 1182-4Wb cells were maintained in a Shields and Sang M3 insect medium (Sigma) supplemented with 10% decomplemented fetal bovine serum and were passaged twice a week at a 1/4 dilution. S2R+ and S2R+ *Wb* cells were maintained in a Schneider insect medium (Dominique Dutscher) supplemented with 10% decomplemented fetal bovine serum and were passaged twice a week at a 1/2 dilution. Cell lines were kept at 25°C.

### Extraction of Wolbachia from cell cultures

The content of ten 25 cm^2^ cell culture flasks reaching confluency with *Wolbachia-* infected JW18 adherent cells was pooled in two 50 mL Falcon tube and centrifuged at 1200 rpm for 5 minutes at room temperature. Next, each pellet was resuspended by pipetting on ice with 3 ml of pre-cooled Nalgene-filtered extraction buffer (220 mM sucrose, 3.8 mM monopotassium phosphate, 8 mM dipotassium phosphate, and 10 mM magnesium chloride).

Cell suspensions were transferred into two 15 ml Falcon conical tubes on ice containing 2 g of sterile 3 mm-glass beads and vortexed vigorously 3 times for 30 seconds with a 30-second incubation period on ice between each round of vortexing.

Each lysate was transferred to a new 15 ml Falcon tube on ice and centrifuged at 1200 rpm for 5 minutes at 4°C. Then, the *Wolbachia-containing* supernatant was transferred to 1.5 mL Eppendorf tubes and centrifuged at 10 000 rpm for 10 minutes at 4°C to pellet *Wolbachia.*

The bacterial pellet of one of the Eppendorf tubes was resuspended in 500 μL of cell culture medium and its content transferred from one tube to another in order to resuspend all the bacterial pellets and collect them in one final tube.

### Generation of the Wolbachia-infected 1182-4Wb and S2R+Wb cell lines

An extract of *Wolbachia* was transferred into a 25 cm^2^ cell culture flask containing confluent 1182-4 or S2R+ cells in a 4 mL volume of cell culture medium. After two days cells were passaged twice a week for a 1-month duration and then, the infection process was repeated to obtain stably infected 1182-4Wb and S2R+Wb cell lines.

To follow the infection dynamics, cells were plated on 18 mm x 18 mm coverslips in a plastic 6-well cell culture plate, and after adherence were fixed in PBS with 3.2% paraformaldehyde for 10 minutes at room temperature, washed for 5 minutes with PBS, and incubated for 2 hours at 37°C in the dark with Alexa Fluor 488 phalloidin A12379 (Life technologies) at a 1/50 dilution. After a 5-minute wash with PBS, coverslips were mounted on glass slides using Fluoroshield with DAPI and observed with an inverted laser scanning confocal microscope (SP5-SMD, Leica Microsystems) using a 63x/1.4 HCX PL APO CS oil objective and images taken with a z-stack interval of 0.5 μm.

The viability of 1182-4 versus 1182-4Wb and S2R+ versus S2R+ *Wb* was evaluated using an automated cell counter (Countess Invitrogen) relying on a trypan blue (Life Technologies) exclusion method according to the protocol of the manufacturer. The cells were passaged the day before the viability measurements were taken.

### Immunofluorescence studies

Cells were plated on 18 mm x 18 mm coverslips in a 6-well cell culture plate 24 hours before fixation in PBS with 3.2% paraformaldehyde for 10 minutes at room temperature. Next, coverslips were dried and immersed in -20°C pre-cooled methanol and kept for 10 minutes at-20°C. Then, coverslips were dried out from residual methanol at room temperature and incubated in a humid chamber for 10 minutes with PBS, BSA 2%. After a PBS wash, cells were incubated for 2 hours at 37°C with the primary antibody or antibodies, added as a 50 μL drop. Following 3 washes of 5 minutes with PBS 1x, cells were incubated for 2 hours at 37°C with the secondary antibody or antibodies. Then, the cells were washed 3 times; each for 5 minutes with PBS 1x and mounted using fluoroshield with DAPI. All primary antibodies were used at a 1/400 dilution: rabbit polyclonal anti-GM130 antibody ab30637 (Abcam) and rabbit monoclonal anti-K48 linkage polyubiquitin antibody ab140601 (Abcam). Mouse monoclonal anti-FK2 ubiquitin antibody AB120 (LifeSensors). Rabbit monoclonal anti-ubiquitin K11 linkage, clone 2A3/2E6 (Millipore). Mouse monoclonal anti-*Wolbachia* surface protein (BEI resources, NIAID, NIH). Secondary antibodies were used at a 1/500 dilution. Goat anti-mouse IgG antibody coupled to Alexa Fluor 488 ab150117 (Abcam), goat anti-rabbit IgG antibody coupled to Cy3 A10520 (Invitrogen). An inverted laser scanning confocal microscope (SP5-SMD; Leica Microsystems) at a scanning speed of 400 Hz equipped with a 63x/1.4 HCX PL APO CS oil objective was used to take images with a z-stack= 0.5 [μm and in the case the images needed deconvolution (Deconvolution software: Huygens Professional version 18.04), the z-stack= 0.2 [μm.

### Drug treatments

Cells were incubated with tunicamycin (Sigma-Aldrich) at 10 μg/mL for 48 hours [62].

### Live experiments

Cells were plated on concanavalin A-coated glass bottom fluorodishes 48 hours before observation. One batch of the S2R+ cell line was treated with tunicamycin as described above. To stain the ER, the cell culture medium was aspirated, cells washed with PBS 1x and incubated for 30 minutes at 25°C with 1 μM live ER-tracker red dye (Molecular Probes) diluted in PBS. The ER-tracker solution was replaced by a 1/20 000 solution of SYTO-11 (Molecular Probes) DNA dye for 10 minutes at 25°C diluted in the appropriate cell culture medium prior to confocal microscopy observations. The temperature of the microscope chamber was set at 25°C prior to observation. For concomitant stainings of the ER and the Golgi apparatus, cells were first incubated for 30 minutes at 4°C with 5 μM of BODIPY FL C_5_-ceramide (Molecular Probes) in PBS. Next, the cells were rinsed 3 times for 2 minutes and incubated for 30 minutes at 25°C with the live ER-tracker red dye as described above. For SP5 confocal time-lapse recordings, stacks of three images, z=0.5p.m, were taken each 5 seconds, with a line average =8, in bidirectional, resonance mode with a SP5 confocal microscope.

To monitor ATF4 activity, cells were plated on concanavalin A-coated glass bottom fluorodishes. Upon cell adherence, the cells were transfected with a 4E-BP intron-dsRed reporter plasmid [43] using the lipofectamine kit (Invitrogen) according to the instructions of the manufacturer. Twenty-four hours post-transfection, one of the fluorodishes containing *Wolbachia-free* cells was treated with tunicamycin (10 μg/ml for 48 hours).

### Image analyses

The image analysis software used is ImageJ version 1.48. The ImageJ macros were developed in collaboration with the MRI-CRBM-Optics plateform, Montpellier, France and are available upon request. The graphing software used was GraphPad Prism version 7.00.

### Electron microscopy

For each cell line, the content of a 25 cm^2^ flask at cell confluence, three days after medium change, was washed and transferred to a 1.5 mL Eppendorf tube and centrifuged at 2000 rpm for 2 minutes at room temperature. The cell pellet was fixed for 1 hour by resuspension in a 2.5% gluteraldehyde-PHEM solution pH=7.4. Fixed cells were kept overnight at 4°C. Cells were next rinced in PHEM buffer and post-fixed in 0.5% osmic acid for 2 hours at room temperature in the dark. After two PHEM washes, cells were dehydrated in a graded series of ethanol solutions (30-100%) before being embedded in EmBed 812 using an automated microwave tissue processor for electron microscopy (Leica AMW). Thin sections of 70 nm were collected at different levels of each block using the Ultracut E microtome (Leica-Reichert). These sections were counterstained with uranyl acetate and lead citrate and observed using a transmission electron microscope (Tecnai F20) at 200 kV.

### RT-qPCR experiments

RNA extraction was performed in biological triplicates for each sample. Precisely, the RNA was extracted from confluent flasks of 25 cm^2^ containing approximately 10^6^ cells. The culture medium was aspirated and replaced by 1 ml PBS 1x. Cells were scraped and transferred to 1.5 ml Eppendorf tubes and centrifuged at 1200 rpm for 5 minutes. Following that, the supernatant was discarded and the cells were resuspended in 300 [μl of the Quick-RNA MicroPrep kit (Zymo Research) lysis buffer. The next steps were performed according to the RNA purification protocol detailed in the kits’ instructions but the in-column DNaseI treatment step was omitted and replaced with a TURBO DNase (Ambion) treatment. RNA was purified using the RNA Clean & Concentrator-5 kit (Zymo Research). cDNA was produced from 2 [μg of RNA using the Superscript VILO cDNA synthesis kit (Invitrogen) and diluted at 1/25 for the RT-qPCR experiments.

Primer pairs were selected according to Primer3 version 0.4.0, synthesized by Eurofins Genomics (S1 table). Primer pairs with an efficiency close to 100% were selected for qPCR experiments.

RT-qPCR reactions were performed using SYBR Green 10x with Platinum Taq (Invitrogen). Amplifications were performed using a Mx3000P instrument (Agilent Technologies) and the MxPro QPCR Software (Agilent Technologies).

The RT-qPCR cycling program consists of a pre-amplification cycle of 2 minutes at 94°C followed by 40 amplification cycles of 30 seconds at 94°C, 30 seconds at 55°C, and 20 seconds at 72°C. The RT-qPCR cycle ends with a dissociation/melt cycle of 1 minute at 94°C, 30 seconds at 55°C, and 30 seconds at 94°C.

For each gene, RT-qPCR is performed in technical and biological triplicates.

The changes in expression were calculated according to the 2^-ΔΔCt^ method [63] and were plotted using the GraphPad Prism software version 7.0.

## Acknowledgments

We thank Alain Debec for critical reading and providing the 1182-4 cell line, William Sullivan for providing the JW18 cell line, and the imaging facility MRI, member of the national infrastructure France-BioImaging supported by the French National Research Agency (ANR-10-INBS-04, investments for the future»), for developing macros for imaging quantification. The ATF4 activity reporter is a kind gift of Hyung Don Ryoo.

## Supporting information

**S1 Fig. Infection of naive cell lines**

(A) Infection dynamics of 1182-4 cells challenged with purified wMel *Wolbachia.* Scoring of intracellular *Wb* was performed on confocal images of fixed cells at the various time points represented on the graph, with a phalloidin staining-yellow-to visualize the cortical actin in order to count the number of intracellular *Wb* only, per individual cells. *Wb* are detected as DAPI bright cytoplasmic foci(-magenta-, i.e. green arrow pointing at a single bacterium at an early time point). Scale bar= 10 [im, n=100 cells per time point, counted in randomly acquired images per coverslip. (B) Cell survival established with Trypan blue. Analyses were performed 24 hr-post medium change.

**S2 Fig. Anti-K48-linkage polyubiquitin immunostainings.**

Confocal acquisitions of the infected and non-infected 1182-4 and S2R+ cell lines stained with WSP-magenta-and an anti-K48-linkage polyubiquitin-yellow-.

**S1 Movie. A *Wolbachia-infected* 1182-4 cell.**

A 1182-4 cell infected by JW18-derived wMel. This animation shows the different Z stacks composing the corresponding confocal merged image in Fig.1. WSP decorates the *Wolbachia* in Cyan and DAPI is in magenta.

**S2 Movie. A *Wolbachia-infected* S2R+ cell.**

A S2R+ cell infected by JW18-derived *wMel.* This animation shows the different Z stacks composing the corresponding confocal merged image in Fig.1. WSP decorates the *Wolbachia* in Cyan and DAPI is in magenta.

**S3 Movie. Time lapse recording of *Wolbachia* and the ER in a 1182-4 *Wb* cell.**

Time lapse acquisitions of a surface focal place in an 1182-4 *Wb* cell. Images are taken each 5 seconds, and the cell is stained with the live DNA dye SYTO 11 to track the *Wolbachia*-magenta-and the ER-tracker is in cyan.

**S1 Table. List of selected primers for qPCR analyses.**

